# Curtailed CaSR expression in peripheral blood mononuclear cells is associated with hypertension

**DOI:** 10.1101/2024.01.23.576550

**Authors:** Sarbashri Bank, Kunal Sarkar, Arindam Chatterjee, Anwesha Das, Santanu Chakraborty, Srilagna Chatterjee, Subhajit Malakar, Dilip Kumar Pal, Sanmoy Karmakar, Sagar Mukherjee, Subrata Kumar De, Somasundaram Arumugam, Madhusudan Das

## Abstract

The calcium-sensing receptor (CaSR) is a GPCR family member. CaSR is abundantly expressed in the parathyroid gland and controls calcium homeostasis. There are no particular discoveries involving CaSR expression in hypertensive peripheral blood mononuclear cells. We investigated lower CaSR expression in hypertensive human and animal models. Nitric oxide (NO) has an important role in vascular function. Impaired NO production disrupts CaSR-mediated fine adjustment of the intra- and extracellular [Ca^2+^] gradients. According to the law Ca^2+^ transportation through Ca channels maintains its connection from the circulatory micro-vascular bed to interstitial space, influenced by capillary and interstitial fluid hydrostatic pressure, as well as oncotic pressure. In that situation, the alterations in Ca^2+^ microenvironment and dynamics remain unknown. CaSR on circulatory mononuclear cells may detect changes in the Ca^2+^ microenvironment in the ISF of endothelial cells-VSMC and endothelium lining in hypertension. In this experiment, the decreased expression of CaSR revealed a molecular sensor in the presence of elevated blood pressure.

---

Calcium sensing receptor (CaSR) is a class C GPCR family member with a large extracellular domain and active dimerization (1). CaSR is highly expressed in the parathyroid (PTH) gland and helps in regulating secretion of parathyroid hormone (2). The extracellular concentration of calcium is monitored by PTH, and thus calcium homeostasis is maintained by the calcitropic action of CaSR in the parathyroid gland and kidneys (3). Although CaSRs have typically been found to be involved in calcium homeostasis, they also play a role in vascular disease and vascular tone modulation (4). However, no specific findings exist regarding CaSR expression in hypertensive peripheral blood-mononuclear cells (PBMCs). We investigated CaSR expression in hypertensive human and rodent, revealing changes in CaSR expression in hypertensive condition.

The study included 55 people with high blood pressure (>140/90 mmHg) along with 35 age and sex-matched healthy controls according to Helsinki declaration and ethical approval (No. IPGME&R/IEC/2021/391). Patients with life-threatening disorders or TSH deficiencies were excluded. Wistar rats (age-sex matched, wt.120 gm, excluding over wt, aged and disease) were included as hypertensive model, administered L-NAME (N^G^-nitro-L-arginine methyl ester) (1mg/mL), a NOS inhibitor (5), for 12 weeks. Non-invasive blood pressure (NIBP) method was applied for the measurement of hypertension in rats. PBMCs were isolated by using Histopaque^®^-1077 in both human and animal studies.

To demonstrate the expression of CaSR in the circulating PBMCs, immunoblotting was performed on PBMC proteins from both patients and animal models. The result showed a reduced CaSR expression in PBMC of hypertensive patients than healthy controls (P<0.05) (Fig-1 A, B, C, D). Similarly, reduced CaSR expression was also observed in hypertensive rat compared controls (P<0.05) (fig. 1 G, H, I, J).

**Fig-1:**
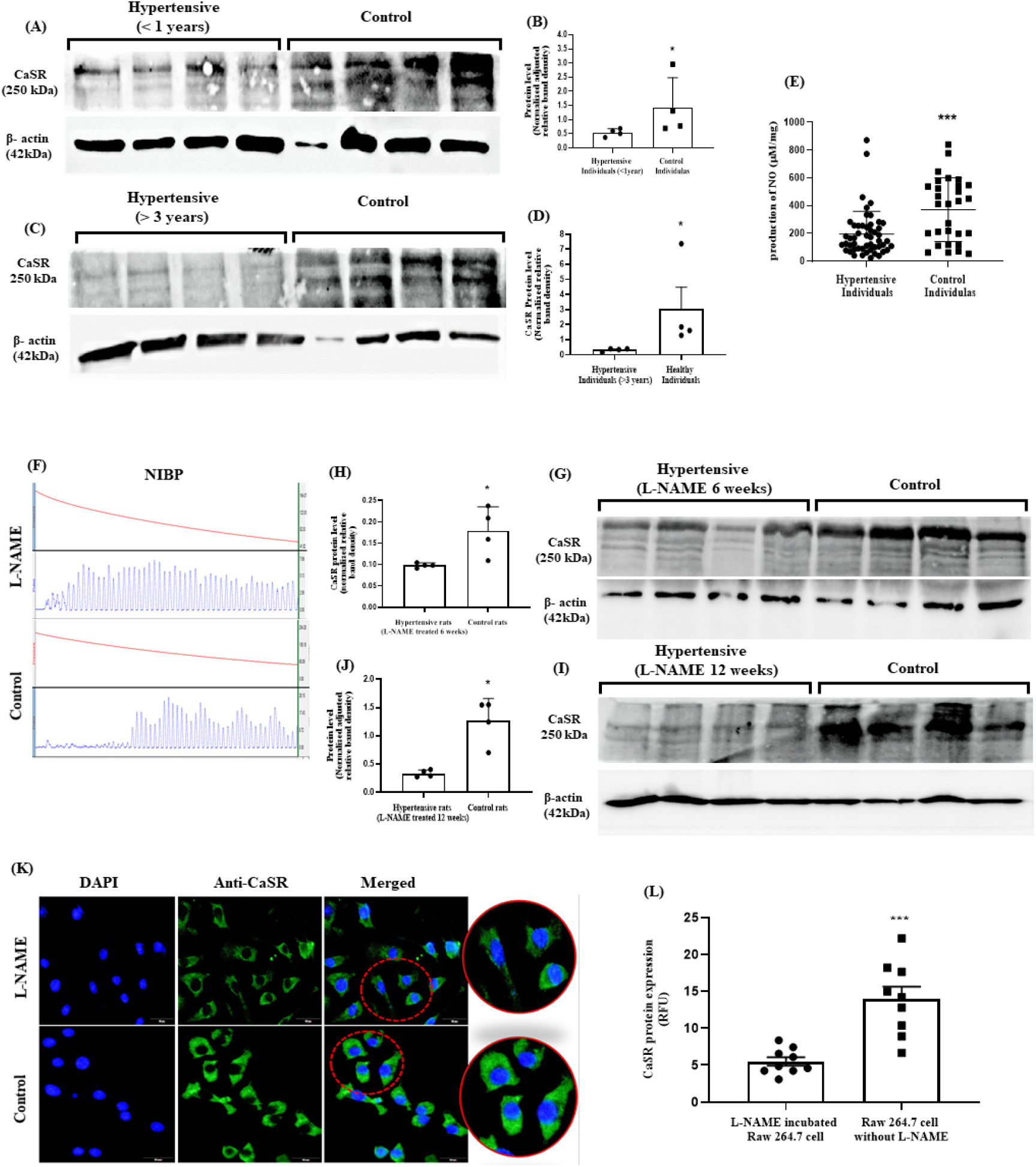
Curtailed CaSR expression (250kDa) in PBMCs of hypertensive subjects: **Fig-1A** demonstrated the expression of CaSR (250 kDa) in PBMC of normal healthy controls (right panel) and hypertensive individuals (left panel) (suffering from hypertension for <1 year), and **fig-1B** showed their difference in CaSR protein level (normalized relative band density) (P <0.05). **Fig-1C, 1D** demonstrated CaSR expression in PBMC of controls and hypertensive patients (suffering from hypertension for > 3 years) and their expression differences (P < 0.05), respectively. **Fig-1E** delineated the nitric oxide level (μM/mg) in hypertensive patients and healthy individuals (P<0.001). The graphs in **Fig-1F** represented the pulse wave of systolic blood pressure in L-NAME (systolic BP: 180±18) treated and control rats (systolic BP: 121±14). **Fig-1G, 1H** represented the CaSR (250 kDa) expression in PBMC of L-NAME treated rats (for 6 weeks) (left panel) and control rats (right panel) and their CaSR level expression differences (P<0.05). **Fig-1I** showed the CaSR expression in L-NAME-treated (12 weeks) hypertensive rats and control rats, and **fig-1J** 1J showed the CaSR level expression difference (P<0.05). **Fig-K** demonstrated the fluorescence expression of CaSR in L-NAME treated raw 264.7 cells (upper panel) and control (no treatment) cells (below panel) [where the left panel showed the DAPI staining, the middle panel showed the fluorescence of anti-CaSR, and the right panel showed a merged DAPI and anti-CaSR (Rabbit mAb-D6D9V)]. **Fig-1L** showed the difference in fluorescence expression (RFU - relative fluorescence unit) of CaSR in L-NAME treated and untreated raw 264.7 cells (P<0.001).

L-NAME treated raw cell line was also used to investigate CaSR expression in mononuclear cells using immunocytochemistry with an anti-CaSR antibody (Rabbit mAb-D6D9V). Under a fluorescence microscope, the expression of CaSR in L-NAME-treated cells was found to be significantly lower (P<0.001) than in untreated cells (Fig-1 K, L).

Nitric oxide (NO) plays a major role in vascular function. As such, the impairment of NO hinders the vasodilation/constriction action. Hence, CaSR expression might be associated with NO generation in hypertensive conditions. To elucidate the link between the generation of NO and CaSR expression, if any, in hypertension, NO was measured in the plasma of each hypertensive and control person by using the Griess method. It was observed that high-blood-pressure individuals produced significantly less NO in their circulation than control individuals (P<0.001) (Fig-1E).

The above study demonstrated that the CaSR expression in PBMCs followed the same paradigm in different experiments in the context of hypertensive models, i.e., we found downregulated CaSR expression in hypertensive PBMCs. Herein, the development of hypertension in an animal model was mediated through the inhibition of nitric oxide synthase (NOS) by using L-NAME. Nitric oxide, the well-known vasodilation agent, plays a major role in vascular diseases. A decrease in NO levels impairs cGMP regulation, causing vascular smooth muscle cell relaxation to be inhibited, resulting in atypical vasoconstriction and increased vascular tone.

Herein, the study categorized hypertensive patients based on their onset of hypertension as described (Fig-1A, C). CaSR expression was found to be downregulated in both conditions, but some discrepancies may occur due to antihypertensive medications impairing absolute CaSR expression. This issue may arise from nitric oxide generation from anti-CVD drugs like aspirin and isosorbide. However, curtailed CaSR expression was detected in hypertensive cases.

On endothelial cells, extracellular [Ca^2+^]_ex_ mediated CaSR activation initiates the signaling of IP_3_-induced Ca^2+^ influx from the sarco-endoplasmic reticulum, and subsequent [Ca^2+^]_cyt_ induced NO generation plays a critical role in hyperpolarization by activating intermediate conductance calcium-activated K^+^ (IKCa) channels. The generated NO then perfuse the VSMC and aids in the synthesis of sGC-activated cGMP, leading to PKG activation, which triggers hyperpolarization by activating big (BKCa) channels in the VSMC (6).

These events downregulate the [Ca^2+^]_cyt_ level by inhibiting voltage-gated Ca^2+^ channels (CaV) and promote the reuptake of Ca^2+^ in the stores and trailing of cytosolic Ca^2+^ extrusion.

As such, decreased Ca^2+^ cues MLCK inactivation, causing smooth muscle cell relaxation. This CaV are very swift Ca^2+^ signalling proteins can mediate a substantial change in a cell. These channels regulate a million Ca^2+^ ions/second down a huge 20,000 folds gradient (1-3mM [Ca^2+^]_ex_ /100-200nM [Ca^2+^]_cyt_). So, thousands of channels/cell can highly increase the Ca^2+^ level within a millisecond (7). Hence, mutilated nitric oxide production in hypertensive patients or NAME induced hypertensive rats may affect the intracellular Ca^2+^ augmentation from the internal storage pool or VDCC activation.

Events can be delineated – Ca^2+^ signalling maintains the autocrine manner during CaSR induced [Ca^2+^]_cyt_ oscillation. So, during [Ca^2+^]_cyt_ spiking, vibrational exportation (by PMCA) of Ca^2+^ can mechanistically stimulate CaSR on the same or paracrine mode. Interestingly, this Ca^2+^ exportation after [Ca^2+^]_cyt_ spiking might promote an upcoming spike by CaSR stimulation on the cell surface. It has also been reported that [Ca^2+^]_ex_ oscillation is the signal that Ca^2+^ cycling actually fortified [Ca^2+^]_cyt_ spiking. So, transient gradients of [Ca^2+^]_ex_ can be initiated due to the spatial and/or temporal difference in efflux and influx of Ca^2+^ (8), where impaired NO induced CaSR modulation can hinder this Ca^2+^ cycling.

It was found that L-NAME acts like a competitive inhibitor of NOS3 enzyme through the binding at substrate (Arg) binding site (356B, 357B, 361B, 366B) of the enzyme. Other co-factor binding site like BH4, FAD, CaM-Ca^2+^. So, downregulated NO

Our key finding that, CaSR expression is downregulated in PBMCs in hypertension, is crucial in the context of NO impairment with fluctuating extra-cellular Ca^2+^ concentration. Impaired NO generation interrupts CaSR mediated fine tuning of intra/extra-cellular [Ca^2+^] gradient. According to the Starling and Kedem-Katchalsky equation, based on trans-vascular fluid flux (Jv) (ml/min) and solute flux (Js) (mg/min) (9), it can be argued that Ca^2+^ transportation through Ca channels maintains its connection from the circulatory micro-vascular bed to interstitial space, influenced by capillary and interstitial fluid hydrostatic pressure and oncotic pressure. In that context, the changes in Ca^2+^ micro-environment and dynamics are still unexplored. Herein, it can be suggested that CaSR on circulatory mononuclear cells can detect changes in the Ca^2+^ microenvironment in the ISF of endothelial cells-VSMC and endothelium lining in hypertension. As such in this experiment the decreased expression of CaSR manifested a significant molecular sensor in case of high blood pressure.

## Acknowledgement

Authors thankfully acknowledge the University Grants Commission for the Fellowship awarded to Sarbashri Bank (D.S. Kothari Postdoctoral Fellowship; BL/20-21/0274). Kunal Sarkar (NET JRF), Anwesha Das (NET JRF), Srilagna Chatterjee (NET JRF).

## References

1. Silve C, Petrel C, Leroy C, Bruel H, Mallet E, Rognan D, Ruat M. Delineating a Ca2+ binding pocket within the venus flytrap module of the human calcium-sensing receptor. J Biol Chem. 2005 Nov 11;280(45):37917–23.

2. Kos CH, Karaplis AC, Peng JB, Hediger MA, Goltzman D, Mohammad KS, Guise TA, Pollak MR. The calcium-sensing receptor is required for normal calcium homeostasis independent of parathyroid hormone. J Clin Invest. 2003 Apr;111(7):1021–8.

3. Chen RA, Goodman WG. Role of the calcium-sensing receptor in parathyroid gland physiology. Am J Physiol Renal Physiol. 2004 Jun;286(6):F1005–11. doi: 10.1152/ajprenal.00013.2004.

4. Schreckenberg R, Schlüter KD. Calcium sensing receptor expression and signalling in cardiovascular physiology and disease. Vascul Pharmacol. 2018 Mar 4:S1537-1891(17)30323-3. doi: 10.1016/j.vph.2018.02.007.

5. Yu J, Wang S, Shi W, Zhou W, Niu Y, Huang S, Zhang Y, Zhang A, Jia Z. Roxadustat prevents Ang II hypertension by targeting angiotensin receptors and eNOS. JCI Insight. 2021 Sep 22;6(18):e133690.

6. Greenberg HZ, Shi J, Jahan KS, Martinucci MC, Gilbert SJ, Vanessa Ho WS, Albert AP. Stimulation of calcium-sensing receptors induces endothelium-dependent vasorelaxations via nitric oxide production and activation of IKCa channels. Vascul Pharmacol. 2016 May;80:75–84.

7. Clapham DE. Calcium signaling. Cell. 2007 Dec 14;131 (6):1047–58. doi: 10.1016/j.cell.2007.11.028. PMID: 18083096.

8. Breitwieser GE. Extracellular calcium as an integrator of tissue function. Int J Biochem Cell Biol. 2008;40(8):1467–80. doi: 10.1016/j.biocel.2008.01.019. Epub 2008 Feb 2. PMID: 18328773; PMCID: PMC2441573.

9. Mehta D, Malik AB. Signaling mechanisms regulating endothelial permeability. Physiol Rev. 2006 Jan;86(1):279–367. doi: 10.1152/physrev.00012.2005. PMID: 16371600.

